# Immunoactive signatures of circulating tRNA- and rRNA-derived RNAs in chronic obstructive pulmonary disease

**DOI:** 10.1101/2024.06.19.599707

**Authors:** Megumi Shigematsu, Takuya Kawamura, Deepak A. Deshpande, Yohei Kirino

**Affiliations:** Computational Medicine Center, Sidney Kimmel Medical College, Thomas Jefferson University, Philadelphia, PA 19107; Center for Translational Medicine, Jane and Leonard Korman Respiratory Institute, Sidney Kimmel Medical College, Thomas Jefferson University, Philadelphia, PA 19107

**Keywords:** short non-coding RNA, circulating RNA, tRNA half, tRNA-derived fragment (tRF), rRNA-derived fragment (rRF), mRNA-derived fragment (mRF), Toll-like receptor 7 (TLR7), plasma, COPD, cytokine

## Abstract

Chronic obstructive pulmonary disease (COPD) is the most prevalent lung disease, and macrophages play a central role in the inflammatory response in COPD. We here report a comprehensive characterization of circulating short non-coding RNAs (sncRNAs) in plasma from patients with COPD. While circulating sncRNAs are increasingly recognized for their regulatory roles and biomarker potential in various diseases, the conventional RNA-seq method cannot fully capture these circulating sncRNAs due to their heterogeneous terminal structures. By pre-treating the plasma RNAs with T4 polynucleotide kinase, which converts all RNAs to those with RNA-seq susceptible ends (5′-phosphate and 3′-hydroxyl), we comprehensively sequenced a wide variety of non-microRNA sncRNAs, such as 5′-tRNA halves containing a 2′,3′-cyclic phosphate. We discovered a remarkable accumulation of the 5′-half derived from tRNA^ValCAC^in plasma from COPD patients, whereas the 5′-tRNA^GlyGCC^half is predominant in healthy donors. Further, the 5′-tRNA^ValCAC^half activates human macrophages via Toll-like receptor 7 and induces cytokine production. Additionally, we identified circulating rRNA-derived fragments that were upregulated in COPD patients and demonstrated their ability to induce cytokine production in macrophages. Our findings provide evidence of circulating, immune-active sncRNAs in patients with COPD, suggesting that they serve as inflammatory mediators in the pathogenesis of COPD.

## Introduction

Chronic obstructive pulmonary disease (COPD) is a progressive respiratory disease, primarily caused by chronic airway inflammation due to exposure to air pollutants, such as smoking and inhalation of environmental chemicals like nitrogen oxides, asbestos, and silicate ^1,2^. While the clinical manifestations of COPD share similarities with those of asthma (e.g., airway hyperresponsiveness, thickened airway walls, and excess mucus secretion), the onset of COPD is typically associated with prolonged exposure to pollutants, leading to a higher prevalence among older individuals ^3, 4^. The main components of COPD are excessive and chronic inflammation, and inflammation-induced remodeling of airways, goblet cell metaplasia, enlargement of mucus-secreting glands, and airway hyperresponsiveness ultimately lead to difficulty in breathing ^5, 6^. Evidence suggests that the host immune response to inhaled stimuli involves activation of macrophages, neutrophils, and leukocytes. In this context, macrophages are key contributors to the pathogenesis of COPD, being involved in the excessive production of various cytokines such as IL-1β, IL-6, IL-8, and TNF-α ^7, 8^. Understanding molecular mechanisms and mediators involved in immune activation in COPD is pivotal for developing effective diagnostic and treatment approaches. In this study, we explored the potential role of a previously unidentified class of inflammatory mediators belonging to extracellular RNAs (exRNAs) in COPD.

Biological processes often take place in extracellular spaces, which encompass biofluids (e.g., blood, cerebrospinal fluid, urine) and extracellular vesicles (EVs). RNA is one of the important components in those extracellular spaces and is referred to as exRNAs or circulating RNAs. In recent years, these exRNAs have garnered increasing attention, partly because of growing recognition that their differential expression patterns serve as potential biomarkers for clinical diagnosis of human diseases including cancers, lung diseases, diabetes, and aging ^9, 10^. Furthermore, the regulatory roles of exRNA have been demonstrated in certain cancers, such that EV-packaged microRNAs (miRNAs) promote cancer metastasis and confer drug resistance ^11–13^. However, the role of exRNAs in the pathogenesis of COPD is not well established.

Recent advancements in RNA sequencing (RNA-seq) and classical biochemical analytics have shown that exRNAs originate from a variety of cellular transcripts, such as messenger RNA (mRNA), miRNA, transfer RNA (tRNA), PIWI-interacting RNA (piRNA), and Y-RNA; also, many exRNAs are encapsulated within EVs, circulating throughout the body ^14–18^. However, standard RNA-seq for short non-coding RNAs (sncRNAs), originally designed for miRNA sequencing, is not suitable for capturing other types of sncRNAs ^19, 20^. The standard approach involves the ligation of 5′- and 3′-adaptors to the 5′-monophosphate (P) and 3′-hydroxyl (OH) ends of sncRNAs. SncRNAs with other terminal forms [i.e., 5′-OH, 3′-P, or 2′,3′-cyclic phosphate (cP)] cannot be ligated with the adaptors, rendering them unamplified and unsequenced. To overcome this limitation, a pre-treatment of RNA samples with T4 polynucleotide kinase (T4 PNK) to leave the ends of all sncRNA species with the 5′-P/3′-OH ends was employed in sncRNA sequencing of human plasma samples ^21–23^ and EVs derived from human macrophages obtained by the differentiation of THP-1 monocyte cells ^24^. The lack of T4 PNK treatment resulted in a dramatic reduction in cDNA yields from EV-sncRNAs ^24^, suggesting that exRNAs predominantly consist of non-miRNAs without the 5′-P/3′-OH ends. These exRNA species have been significantly underrepresented in current sncRNA analyses using standard RNA-seq, thereby representing a largely unexplored field in sncRNA transcriptome research ^19, 20^.

We herein report a comprehensive characterization of circulating sncRNAs in plasma from patients with COPD. Previous investigations of circulating sncRNA expression in COPD patients relied on standard RNA-seq without T4 PNK treatment ^25–28^, thus focusing mainly on miRNAs. By conducting sncRNA sequencing on plasma RNA samples pre-treated with T4 PNK, we have established the expression profile of plasma sncRNAs and their alterations between healthy individuals and COPD patients. Our detailed examination on tRNA-derived sncRNAs revealed abundant and specific accumulation of tRNA^ValCAC^-derived sncRNA in COPD patients. Interestingly, the tRNA^ValCAC^-derived sncRNA activates human macrophages via TLR7 and induces cytokine production. Our studies also identified rRNA-derived sncRNAs that were upregulated in COPD patients and have a potent ability to induce cytokine production in macrophages. Our results provided insights into regulation of circulating sncRNA repertoire and its potential roles as a unique class of inflammatory mediators in the COPD.

## Results

### The levels of 5′-HisGUG are upregulated in plasma from patients with COPD

Our recent study revealed that a specific extracellular tRNA-derived sncRNA [i.e., 5′-tRNA half from tRNA^HisGUG^ (5′-HisGUG)] is delivered into endosomes of human macrophages, possessing a potent ability to stimulate endosomal Toll-like receptor 7 (TLR7) and induce cytokine production ^24^. Given the activity of 5′-HisGUG and considering the role of macrophages and excessive cytokines in COPD pathogenesis ^7, 8^, we first quantified the levels of 5′-HisGUG (**Table S1**) in plasma samples from healthy individuals and patients with COPD. We used our established multiplex version of TaqMan RT-qPCR, which enables simultaneous quantification of a specific tRNA half and an internal control in limited sample quantities ^29^. As a result, the COPD samples showed a significant upregulation of 5′-HisGUG levels [3.75 fold (*p*=0.028, by Student’s *t*-test)] (**Fig. S1**), prompting us to proceed to a comprehensive characterization of all circulating tRNA-derived sncRNA species in plasma from healthy individuals and COPD patients.

Since COPD is an age-dependent disease with higher prevalence in older adults, and considering our previous study suggesting age-dependent changes in tRNA half accumulation in various mouse tissues ^30^, we aimed to minimize the effects of age on sncRNA expression by obtaining plasma samples for sequencing from each four healthy and four COPD donors within a similar age group of 50–70 years old (**Table S2**). We treated RNAs extracted from each plasma sample with T4 PNK in the presence of ATP to convert the RNA ends to 5′-P/3′-OH (**Fig. 1A**), which was followed by adaptor ligation, cDNA amplification, and Illumina sequencing. In cDNA purification and bioinformatic analysis, we targeted sncRNAs ranging from 18 to 50 nucleotides (nt) in length. Principal component analysis (PCA) on tRNA-mapped reads revealed that tRNA-derived sncRNAs in COPD patients form a cluster distinct from that of healthy donors (**Fig. 1B**).

**Figure 1.**
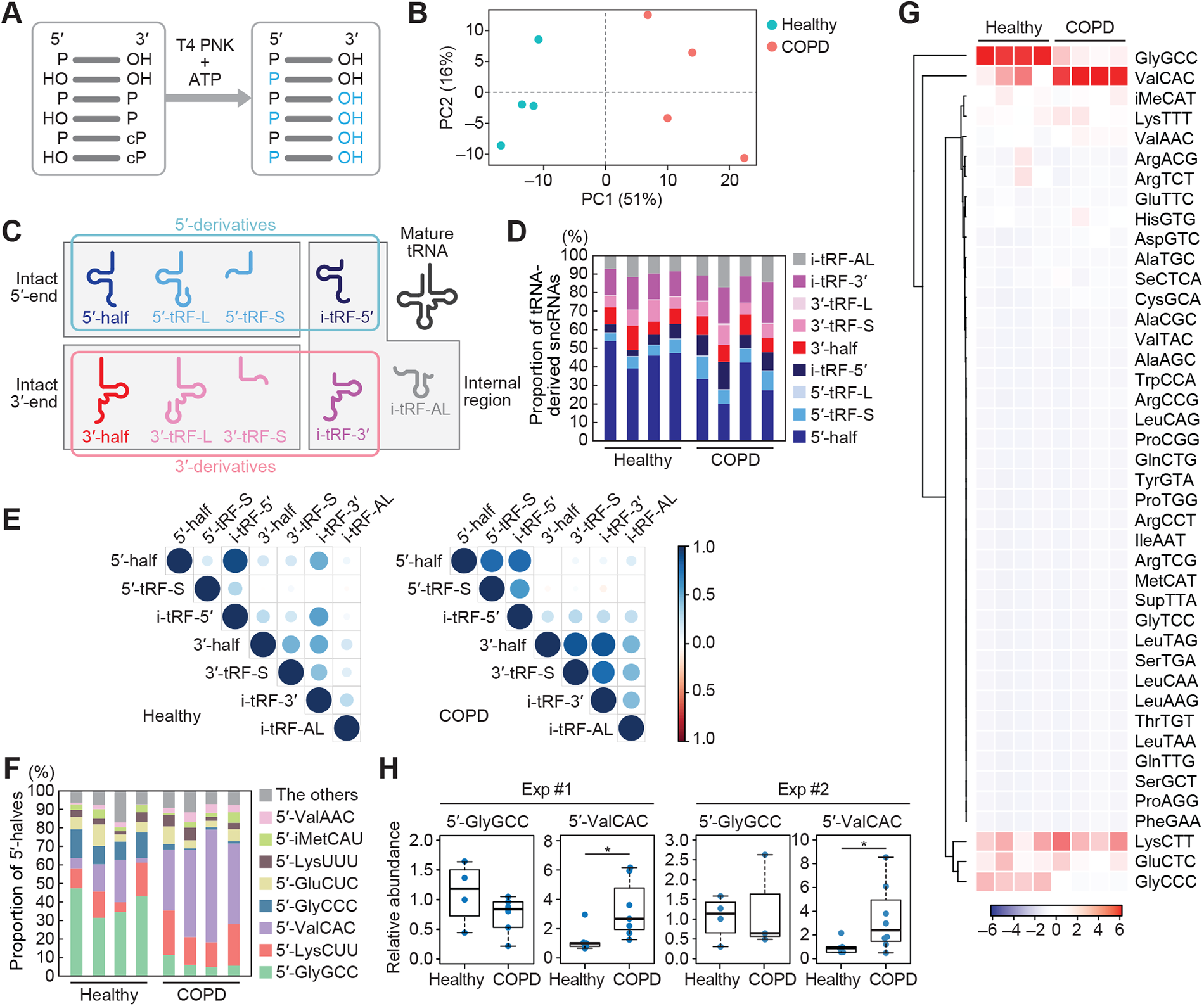
Analysis of tRNA-derived sncRNAs in human plasma samples. **(A)** Schematic representation of the sncRNA sequencing for T4 PNK-treated RNAs. **(B)** PCA of tRNA-mapped reads. **(C)** Classification of tRNA-derived sncRNAs in this study. **(D)** Proportion of tRNA-derived sncRNAs classified to each subclass. **(E)** Spearman correlation analysis among each class of tRNA-derived sncRNAs. The color and size of each dot indicate the correlation coefficient calculated as Z-value. **(F)** Proportion of 5′-halves derived from respective cytoplasmic tRNA isoacceptors. **(G)** Heatmap and dendrogram showing the clustering of the 5′-half expression pattern. **(H)** The levels of 5′-GlyGCC and 5′-ValCAC in plasma samples quantified by TaqMan qPCR. The value of one of the healthy samples was set as 1, and the relative values for the other samples are shown. The sample size in the first set of experiment: Healthy: n = 5; COPD: n = 7. For 5′-GlyGCC, only four healthy samples and six COPD samples were detected and included in the graph. The second set of experiment: Healthy: n = 5; COPD: n = 8. For 5′-GlyGCC, only four healthy samples and three COPD samples were detected. \**p*<0.05.

### 5′-tRNA halves constitute the most abundant tRNA-derived sncRNAs in plasma

For profiling, the identified tRNA-derived sncRNAs were classified based on tRNA fragment (tRF) classification method ^31–33^, incorporating modifications from our previous study ^30^ as illustrated in **Fig. 1C**. A tRNA-derived sncRNA produced by the anticodon cleavage is defined as a tRNA half, which can be further categorized as either the 5′-half or the 3′-half, each retaining their respective intact ends. tRFs encompass any tRNA fragments with the exception of tRNA halves. Note that some reports describe any tRNA fragments, including halves, as tRFs; however, we distinguish between tRNA halves and tRFs. The 5′-tRF retains an intact mature 5′-end and is produced through a 3′-end cleavage occurring outside of the anticodon loop. In this study, it is further classified into two groups: the Short (5′-tRF-S) has a shorter length, resulting from a cleavage upstream of the anticodon loop; the Long (5′-tRF-L) has a longer length, produced from a cleavage occurring downstream of the anticodon loop. Similarly, the 3′-tRF can be classified as 3′-tRF-S or 3′-tRF-L, depending on whether the 5′-end cleavage occurs downstream or upstream of the anticodon loop, respectively. Furthermore, we have classified internal-tRF (i-tRF), which lacks both intact 5′- and 3′-ends, into three subclasses: i-tRF-5′, i-tRF-3′, and i-tRF-AL. The i-tRF-5′ and i-tRF-3′ are derived from the 5′- and 3′-parts of mature tRNAs, respectively, while the remaining subclass, i-tRF-AL, encompasses the entire anticodon loop. We define 5′-half, 5′-tRF, and i-tRF-5′ as 5′-derivatives, and 3′-half, 3′-tRF, and i-tRF-3′ as 3′-derivatives.

In both healthy and COPD plasma, the most predominant class of tRNA-derived sncRNAs was the 5′-half (**Fig. 1D**). Although the proportion of 5′-half appeared smaller in COPD compared to the healthy plasma, this does not necessarily indicate a decrease in the actual abundance of 5′-halves, given that the increased level of 5′-HisGUG was experimentally confirmed (**Fig. S1**). It is important to note that much of the bioinformatic data are presented as relative abundance of different sncRNAs within a sample and does not indicate quantitative difference between samples. We further investigated quantitative relationship among each subclass by *Spearman* correlation analysis (**Fig. 1E**). In the healthy cohort, a strong correlation was observed only between 5′-half and i-tRF-5′. In COPD, not only i-tRF-5′ but also 5′-tRF-S showed a significant correlation with the 5′-half, while i-tRF-3′ and 3′-tRF-S exhibited a correlation with the 3′-half. These findings may indicate that the processing of tRNA halves to the tRF-S and i-tRFs is enhanced in COPD. There was an increased proportion of i-tRF-5’ and 5′-tRF-S, whereas the 5’-half showed a decreased proportion in COPD (**Fig. 1D**).

### COPD patients exhibit differential profiles of 5′-halves compared to healthy individuals, with 5′-ValCAC being the most predominant tRNA-derived sncRNAs

Since the 5′-half is the most abundant class of tRNA-derived sncRNAs and 5′-HisGUG is upregulated in COPD patients, we further analyzed the constituent species of 5′-half accumulated in human plasma samples. Although human tRNAs have more than 40 isoacceptors, the primary tRNA-derived sncRNAs found in human plasma were three 5′-haves derived from tRNA^GlyGCC^, tRNA^ValCAC^, and tRNA^LysCUU^, collectively constituting 60.5–79.3% of 5′-halves (**Fig. 1F**) and 13.5–34.5% of tRNA-derived sncRNAs (**Fig. S2**). While 5′-GlyGCC was identified as the most abundant 5′-half in healthy individuals, 5′-ValCAC exhibited the highest abundance in COPD patients (**Fig. 1F and 1G**). Their cognate isoacceptors exhibited a similar trend; the relative abundance of 5′-GlyCCC decreased while that of 5′-ValAAC increased in the COPD compared to healthy samples. These differential profiles were observed in all eight of the plasma samples used in this study, indicating their significant consistency. The second most abundant species was 5′-LysUCC in both groups. While our TaqMan RT-qPCR quantification showed a modest decrease of 5′-GlyGCC in COPD samples, the levels of 5′-ValCAC were higher [3.38 fold (*p* = 0.029) and 3.34 fold (*p* = 0.048) in two independent analyses, respectively; **Fig. 1H**], replicating the sequencing results and thus experimentally validating the upregulation of the levels of 5′-ValCAC in COPD patients. These findings suggest that the production of specific 5’-halves is selectively and consistently promoted in COPD.

Concerning differences between healthy and COPD samples in tRNA-derived sncRNAs, we further noticed distinctions in the 3’-terminal region of 3’-halves. The mature tRNA molecule has a strictly conserved trinucleotide sequence at the 3′-end, cytosine-cytosine-adenosine (CCA), which serves as the attachment site for amino acids (**Fig. S3A**). Previous studies from cP-RNA-seq, which specifically sequences sncRNAs with a cP ^34, 35^, indicated a significant deficiency of the terminal A nucleotide of CCA in 3′-halves across human cell lines as well as mouse tissues ^30,36^. The lack of the terminal A is also observed in mature tRNA, though in small proportions ^37^. Here, sequencing of T4 PNK-treated sncRNAs enabled us to accurately detect the proportion of 3′-terminal variants in all circulating 3′-halves in human plasma samples, regardless of their terminal phosphate forms (**Fig. S3B**). In the healthy group, 3′-halves lacking the 3′-terminal CA sequence were the most abundant. In COPD samples, 3′-halves with an intact CCA end were the most abundant, followed by 3′-halves lacking the 3′-terminal A nucleotide. These results suggest that the anticodon cleavage of mature tRNAs is enhanced in COPD, potentially leading to a higher proportion of 3′-halves with an intact CCA-or CC-end.

### The primary tRNA halves are produced from specific isodecoders

We further analyzed the expression profiles of tRNA-derived sncRNAs, focusing specifically on those derived from three major tRNA sources: tRNA^GlyGCC^, tRNA^ValCAC^, and tRNA^LysCUU^. Consistent with the proportion of total tRNA-derived sncRNAs (**Fig. 1D**), the 5′-half was the most abundant species among the sncRNAs derived from each tRNA, in both healthy and COPD samples (**Fig. 2A**). The source of the 5′-half was analyzed based on isodecoders, unique sequences of tRNA distinguished by the gene ID ^38^ (**Fig. 2B, Fig. S4A**). Despite the human genome encoding 5 isodecoders for tRNA^GlyGCC^, 5′-GlyGCC was exclusively derived from three genes, GlyGCC-2-1, 3-1, and 5-1 (**Fig. 2B**). The sequences of 5′-GlyGCC-2-1 and 3-1 are identical, while their 3′-halves differ by one nucleotide (**Fig. S4A**). Because 3′-GlyGCC was hardly detected (**Fig. S2**), it is impractical to determine which isodecoder, GlyGCC-2-1 or 3-1, truly serves as the source of 5′-GlyGCC. For tRNA^ValCAC^, 3 out of the 6 isodecoders encoded in the human genome contributed to the production of 5′-ValCAC, and the sequences of these three 5′-halves are identical (**Fig. S4A**). Further, only three out of the 10 isodecoders contributed to the production of 5′-LysCUU. There was no difference in the detected isodecoders for the 5′-halves between the healthy and COPD samples, which could be due to identical profiles of the expressed isodecoders and/or the isodecoders subjected to anticodon cleavage.

**Figure 2.**
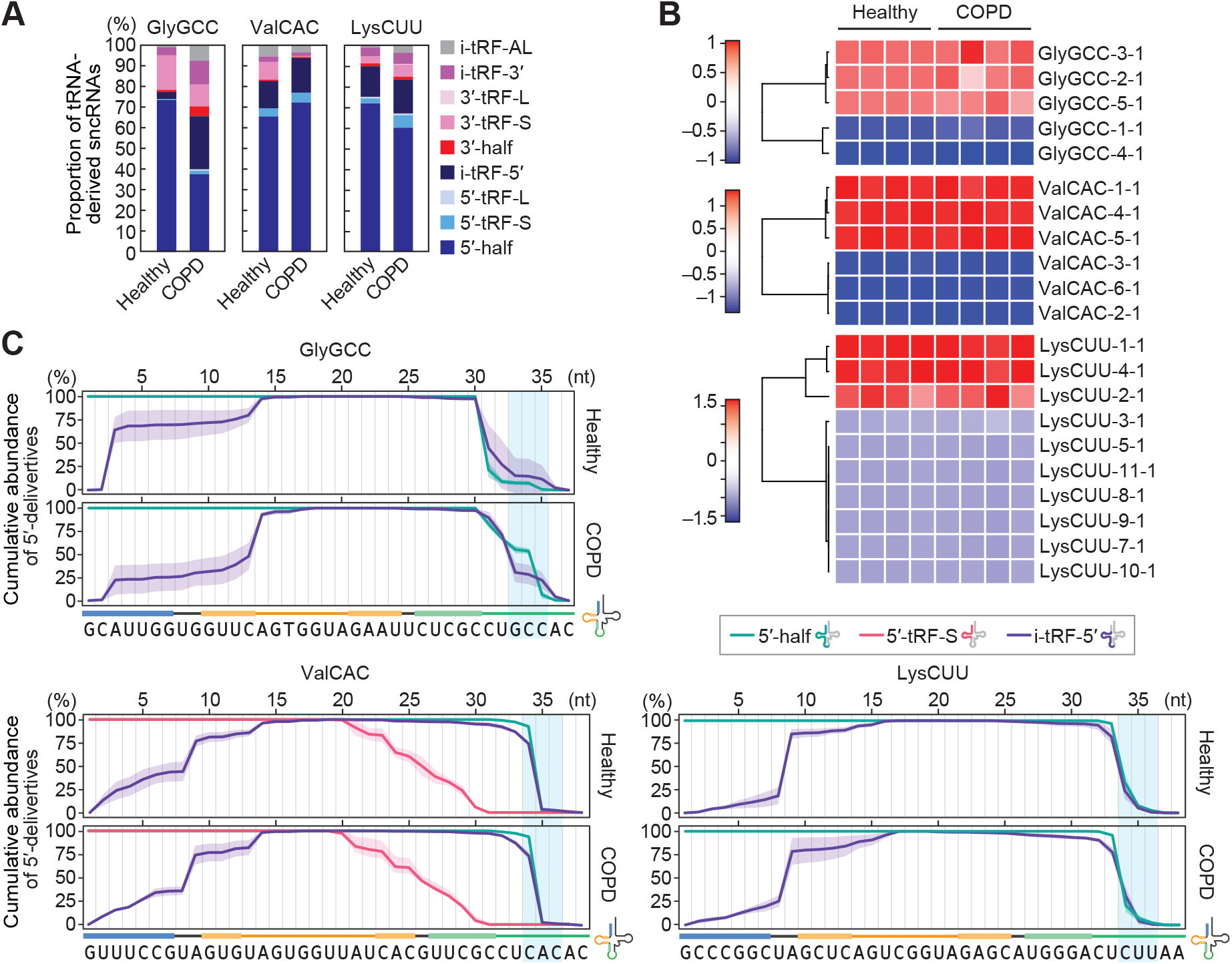
Analysis of the sncRNAs derived from tRNAGlyGCC, tRNAValCAC, and tRNALysCUU. **(A)** Proportion of the reads derived from the indicated three tRNAs, classified into each subclass of tRNA-derived sncRNAs. **(B)** Heatmap of the expression of sncRNAs derived from each isodecoder. **(C)** Analysis of read counts for each classified tRNA-derived sncRNA. The highest read count in each subclass of tRNA-derived sncRNAs was set as 100%, and the read counts at each nucleotide position are shown as relative abundances. The shaded region overlaying each cumulative line represents the standard deviation (SD) from four replicates.

The sequence reads derived from the three most abundant isodecoders for each of the three tRNAs were further analyzed at single nucleotide level. In the data of tRNA^GlyGCC^-derived reads, read counts of 5′-half and i-tRF-5′ are displayed as a cumulative line (**Fig. 2C**). The 3′-end of 5′-GlyGCC is not consistent, appearing to vary in both healthy and COPD samples. The 3′-teminal of the 5′-half and the i-tRF-5′ did not align well, suggesting that i-tRF-5′-GlyGCC might not be directly produced from 5′-GlyGCC. The cumulative line of i-tRF-5′-GlyGCC indicates two sites where i-tRF-5′-GlyGCC is produced, between C2 and A3 in the acceptor stem and between C13 and A14 in the D-stem-loop. Unlike tRNA^GlyGCC^-derived sncRNAs, almost all 5′-ValCAC and i-tRF-5′-ValCAC sequences have a length of 34 nt, ending at C34, which is the first nucleotide of the anticodon (**Fig. 2C**). 5′-tRF-S-ValCAC, which showed abundant read counts and was also plotted, exhibited gradual excision from the 3′-end. In contrast, i-tRF-5′-ValCAC showed a singular endo-excision site between U8 and A9. Similar to tRNA^ValCAC^-derived sncRNAs, the 3′-end of 5′-LysCUU was consistent with that of the i-tRF-5′. The i-tRF-5′-LysCUU also had endo-excision between U8 and A9. These findings collectively suggest that the production of tRNA-derived sncRNAs follows a distinct mode of action depending on the tRNA species. In the cases of tRNA^ValCAC^ and tRNA^LysCUU^, i-tRF-5′ is predicted to be produced from the cognate 5′-half, a phenomenon not observed in tRNA^GlyGCC^. The production of i-tRF-5′ tends to occur through cleavage at specific sites, particularly between C and A, or U and A nucleotides, indicating the involvement of specific endoribonucleases.

In addition to the three tRNAs, we also analyzed the reads derived from tRNA^HisGUG^. tRNA^HisGUG^ is unique in that it contains an extra guanosine at the 5′-end, referred to as G_–1_, which is added post-transcriptionally by a specific guanylystransferase, THG1 ^39^. G_–1_ of tRNA^HisGUG^ is highly conserved and plays a crucial role in aminoacylation, as it is recognized by histidyl-tRNA synthetase (HisRS). Our previous study demonstrated that human cell lines contain a fraction of mature tRNA^HisGUG^ molecules lacking G_–1_, and that a substantial portion of the 5′-HisGUG lacks G_–1_ and instead has G_+1_ as the 5′-end in human cell lines and mouse tissues^24, 30, 36, 40^. This study is the first to report the proportion of G_–1_ and G_+1_ in 5′-HisGUG in human plasma samples. The majority of tRNA^HisGUG^ were mapped to the isodecoder HisGUG-1-1 (94.2%, **Fig. S4A and S4B**). The analyses of 5′-HisGUG sequences mapped to HisGUG-1-1 revealed that over 70% of them lack G_–1_ (**Fig. S4C and S4D**). The predominant 3′-end of 5′-HisGUG was G34, consistent with our previous findings in human cell lines and mouse tissues ^24, 30, 36, 40^.

### 5′-ValCAC induces cytokine production via TLR7 activation in macrophages

In the development of COPD, macrophages act as important mediators of inflammation by secreting cytokines in response to exposure to harmful substances ^7, 8^. We recently reported that 5′-HisGUG can induce cytokine production by serving as a potent endogenous ligand of endosomal TLR7 in human macrophages derived from THP-1 ^24^. In our recent broader characterizations of various 5′-tRNA halves, we further identified 5′-ValCAC as another active molecules in TLR7 activation ^41^. The 5 -ValCAC found in EVs secreted from human macrophage cells was 33 nt, one nucleotide shorter than the most abundantly found 34-nt version in human plasma. Given that the 34-nt version of 5′-ValCAC was the most predominant and upregulated in COPD patients (**Fig. 1F-1H, 2A, 3A**), we examined its activity in simulating the macrophage cells and inducing the production of cytokines such as TNFα, IL-1β, and IL-6. 5′-ValCAC was delivered into endosomes of macrophages using a cationic liposome reagent, DOTAP, which mimics exosome and has been widely used for endosomal delivery in previous studies ^24, 41–44^. For controls, a 20-nt HIV-1-derived ssRNA termed ssRNA40, known to strongly activate endosomal TLRs ^45^, and its inactive mutant (ssRNA41), in which U is replaced with A, were used. Remarkably, the levels of all three examined cytokines secreted into the culture medium significantly increased upon endosomal delivery of 5′-ValCAC (**Fig. 3B**). Notably, the cytokine secretion levels induced by 5′-ValCAC were significantly higher compared to those by ssRNA40. The 3′-ValCAC, in contrast to its counterpart 5′-ValCAC, did not induce cytokine secretion (**Fig. 3B**). We then evaluated whether the cytokine production is mediated by the activation of TLR7 by using *TLR7* knockout (KO) THP-1 cells. Induction of cytokine secretion was not observed in *TLR7* KO cells (**Fig. 3C**), indicating that 5′-ValCAC strongly activates macrophages via TLR7.

**Figure 3.**
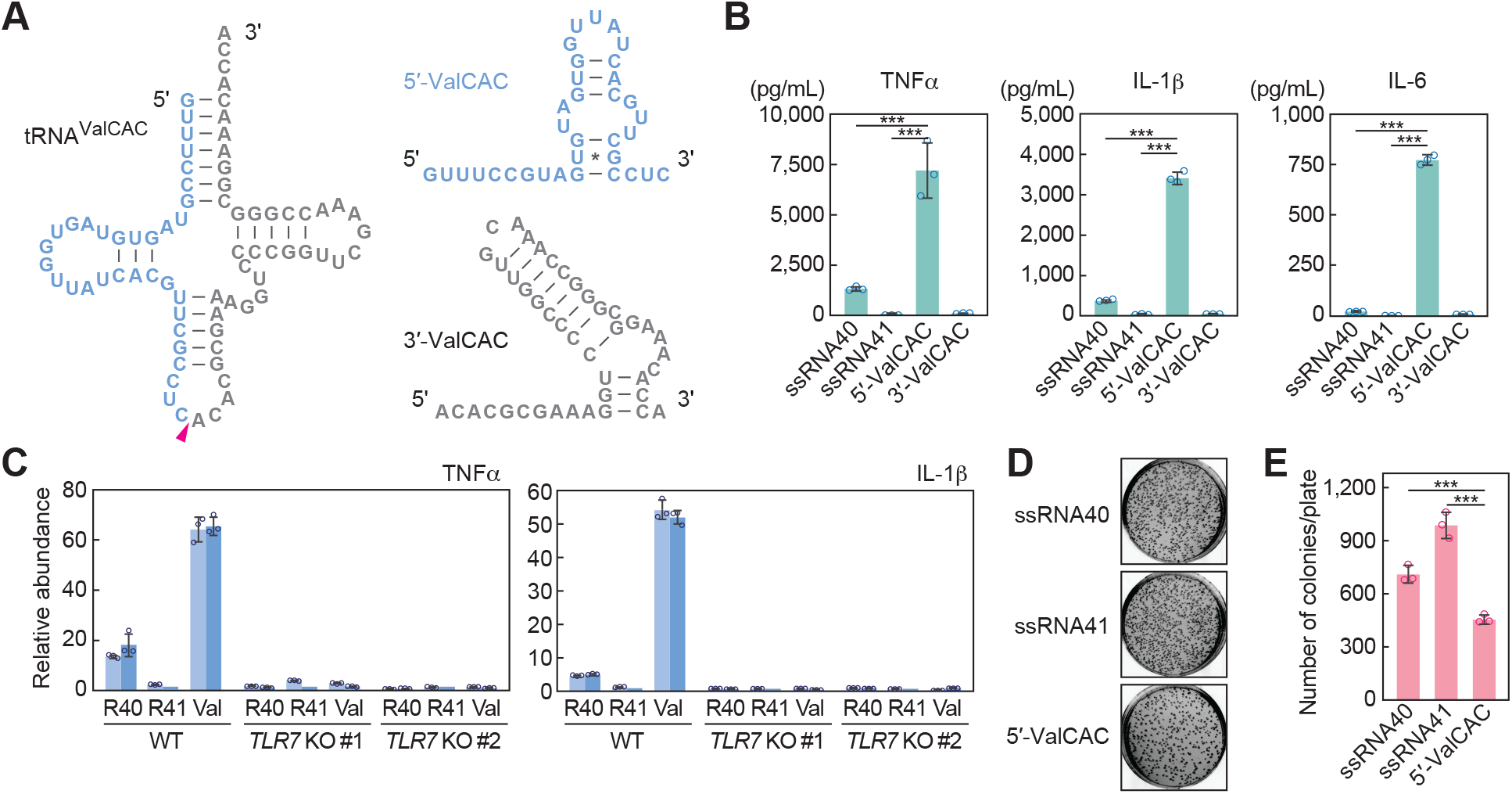
5′-ValCAC activates TLR7 in human macrophages. **(A)** Secondary structure of mature tRNAValCAC, 5′-ValCAC, and 3′-ValCAC (predicted by mFold^46^). **(B)** After DOTAP-mediated endosomal delivery of the indicated RNAs into macrophages, culture medium was subjected to the measurement of concentration of the indicated cytokines. The error bars denote the SD of three independent experiments. \*\*\**p* < 0.001. **(C)** The DOTAP experiments were performed by using two different *TLR7* KO THP-1 cell clones (#1 and #2), followed by quantification of cytokine mRNAs by RT-qPCR. Two independent experiments were performed, which are shown as separated bars, while the error bars indicate the SD of qPCR replicates. **(D, E)** After DOTAP-mediated endosomal delivery of the indicated RNAs, macrophages were subjected to bacterial infection and invasion assay. Representative pictures of the plates with *E.coli* colonies (D) and bar graphs of the counted numbers of colonies (E) are shown. The error bars indicate the SD of three independent experiments. \*\*\**p* < 0.001.

We further investigated whether the 5′-ValCAC-mediated macrophage activation contributes to bacterial elimination. After DOTAP-mediated endosomal delivery of 5′-ValCAC, human macrophage cells were infected with *E.coli*. Following incubation, the *E. coli*-infected macrophages were lysed, and viable *E.coli* were rescued on LB agar plates. As shown in **Fig. 3D-E**, *E. coli* colony numbers were significantly reduced on the plates from macrophages transfected with 5′-ValCAC compared to those transfected with the negative control ssRNA41. ssRNA40 also decreased colony numbers relative to ssRNA41, but the reduction was less pronounced than that by 5′-ValCAC (**Fig. 3D-E**). These results align with the levels of cytokine induction (**Fig. 3C**), and suggest that 5’-ValCAC-mediated activation of macrophages is a fully functional response and may elicit an immune response against secondary infections.

### Differential profiles of plasma rRNA-derived fragments (rRFs) between healthy individuals and COPD patients

Circulating RNAs can originate from transcripts other than tRNAs ^14–18^. It is postulated that long transcripts circulate as processed fragments, and fragments derived from rRNA and mRNA are considered potential biomarkers ^9, 10^. Among plasma samples, rRFs were found to be the most abundant sncRNA species (**Table S3**), prompting us to further analyze the reads of rRFs. PCA exhibited modest differences between healthy and COPD samples (**Fig. 4A**). There appears to be more variation among the healthy samples, while the patient samples tend to converge. Most rRFs are derived from 28S and 18S rRNAs (**Fig. S5A and S5B**). In contrast to our previous cP-RNA-seq analysis of human cell lines and mouse tissues, which displayed distinct peaks in the alignments ^30, 36^, the plasma data in this study showed scattered and modest peaks throughout the transcript (**Fig. S5C**). As suggested by the PCA, modest differences were observed in the alignment between healthy and COPD samples.

**Figure 4.**
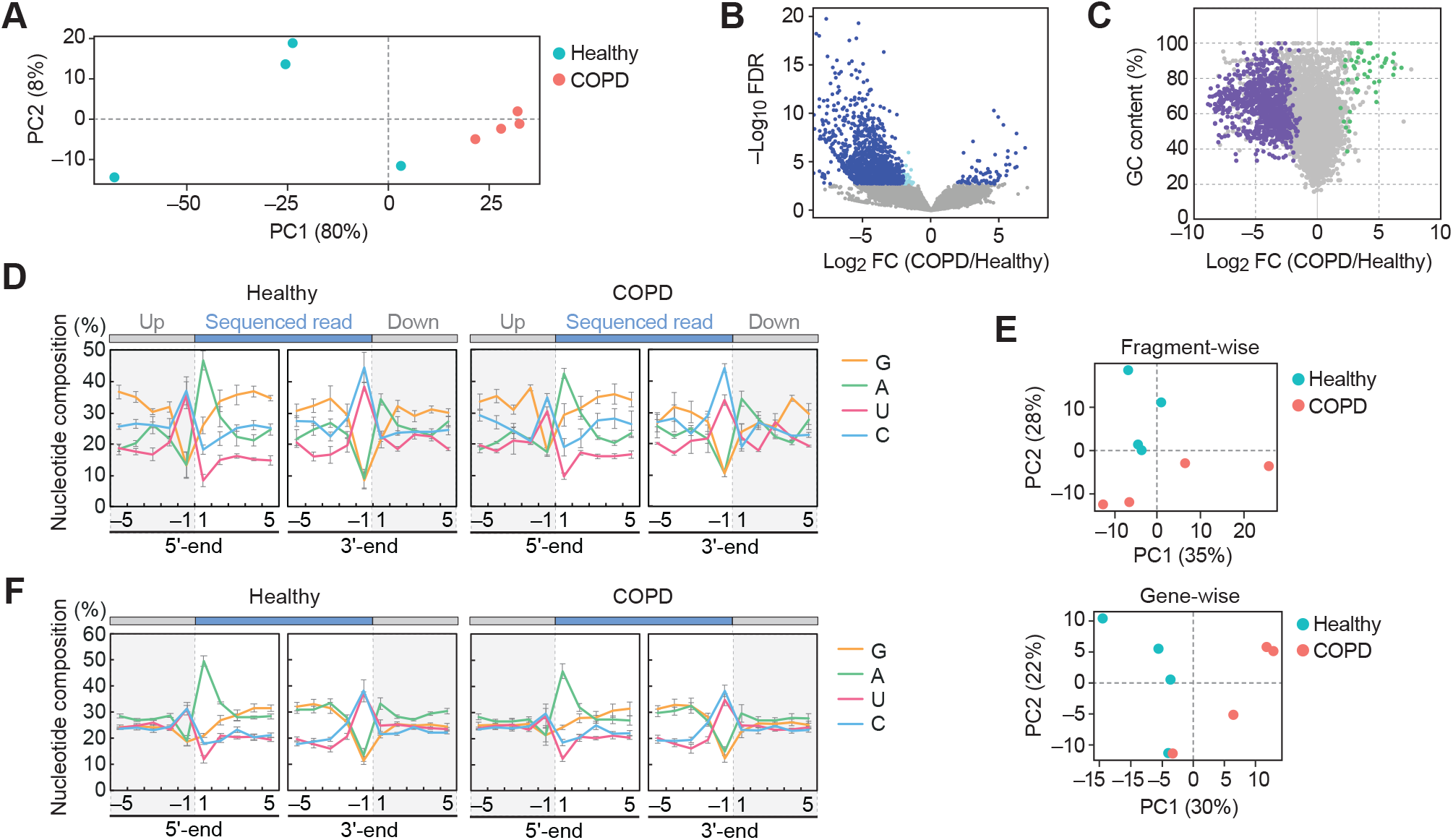
Analysis of plasma rRFs and mRFs. **(A)** PCA of rRFs. **(B)** Volcano plot for rRFs obtained from DESeq2. The light blue dots indicate a fold change level lower than 4-fold or higher than 4-fold, while the navy blue dots represent FDR of less than 0.05. **(C)** The GC contents of each rRF were plotted against their log2 fold change values. The rRFs with an average count among 8 libraries lower than 10 were excluded, and 48 upregulated rRFs and 879 downregulated rRFs with FDR < 0.05 are depicted in green and purple, respectively. **(D)** Nucleotide composition around the 5′- and 3′-ends of the rRFs. This analysis was performed based on the number of reads, thus reflecting the abundance of each sequence read. A dashed line separates upstream and downstream positions for the 5′- and 3′-ends, representing the cleavage site that generates rRFs (the regions outside of rRF are colored in grey). **(E)** PCA of mRFs. PCA was performed using either fragment counts (left panel) or mRNA ID base (right panel). **(F)** Nucleotide composition around the 5′- and 3′-ends of the mRFs.

We performed a differential expression analysis on the rRFs, which unexpectedly revealed a propensity for decreased expression of rRFs in the COPD samples. Among over 40,000 unique rRF species, 1,459 rRFs exhibited a false discovery rate (FDR) lower than 0.05. Within this group, 67 rRFs were upregulated in COPD, while 1,392 rRFs were downregulated (**Fig. 4B**). The rRFs exhibiting the most significant changes, with the lowest false discovery rate (FDR), displayed more than a 20-fold decrease in the COPD samples (**Table 1**). We then aimed to identify upregulated rRFs in COPD and found a 24-nt fragment, derived from nucleotide position (np) 803 of the 28S rRNA, ranking 68th in terms of the lowest FDR. This specific rRF exhibited a high GC content of 91.7%, leading us to hypothesize that upregulated rRFs in COPD may have a high GC content. Indeed, upregulated rRFs displayed higher GC content (average: 79.85 ± 2.41%), while downregulated rRFs displayed lower GC content (average: 63.02 ± 1.61%) (**Fig. 4C**). These findings imply that rRFs are more susceptible to cleavages by ribonucleases in COPD, resulting in the upregulation of rRFs with high GC content being the predominant species. We also analyzed the nucleotide composition of the rRFs, including both upstream and downstream regions. In these analyses, we excluded the read starting from np3303 in 28S rRNA (**Fig. S5C**), whose extensive abundance can have a biased impact on the composition. As shown in **Fig. 4D**, the terminal end of rRFs predominantly exhibits an aggregation of A at the 5′-end, while C or U are more prevalent at the 3′-end. This pattern is consistently observed in both the upstream and downstream regions, suggesting the involvement of specific endoribonucleases that tend to cleave between C and A or U and A in the production of rRFs.

### Profiles of mRNA-derived sncRNAs in plasma

We next analyzed reads derived from mRNA. In contrast to tRNA and rRNA fragments, mRNA-derived fragments (mRFs) did not show a significant difference between healthy and COPD samples, as observed in both fragment-wise and gene-wise analyses (**Fig. 4E**). We found that mRFs were derived from specific reads (**Fig. S5D**). These fragments could have multiple annotations in mRNA or genomic regions, making it difficult to conclusively determine if they were derived solely from one mRNA. However, mRFs exhibited unique nucleotide compositions, although there were no significant differences between healthy and COPD samples (**Fig. 4F**). Nearly half of the fragments begin with A and terminate with U or C. This suggests that the ribonuclease involved in rRF production may also contribute to the generation of the 3′-end of mRFs, while the 5′-end of mRFs could be produced in a manner distinct from that of rRFs.

### rRFs function as immunoactive molecules by activating endosomal TLR

Next, we sought to determine the potential immunomodulatory role of rRFs upregulated in COPD. We identified a 39-nt rRF starting from np21 of 18S rRNA, which was upregulated 5.8-fold in COPD with an FDR of 0.035. This fragment contains two UU sequences and two GU sequences, which can be a preferable ligand for TLR7 ^45^. Due to low yield after synthesis, we synthesized the RNA starting from np22, instead of np21, hereafter referred to as 18S-np22 (**Fig. 5A**). In addition, we selected another rRF, 28S-np4533, which exhibited 1.35-fold upregulation and contained three UU and four GU sequences (**Fig. 5A**). The predicted secondary structure of these rRF sequences, including 50-nt upstream and downstream regions, revealed that rRFs are located in stem-loop regions (**Fig. 5B**). Interestingly, even without the inclusion of upstream and downstream regions, the rRFs exhibited the same secondary structure, as predicted by mFold ^46^. This suggests that the rRFs adopt a stable conformation independent of their surrounding sequences. We delivered these two rRFs into the endosomes of macrophages to examine macrophage activation potential via endosomal TLR. As a result, both 18S-np22 and 28S-np4533 induced the cytokine production (**Fig. 5C**). As observed in the case of 5′-ValCAC, the cytokine secretion levels induced by 18S-np22 were significantly higher compared to ssRNA40-induced levels, while the cytokine levels induced by 28S-np4533 were comparable to those induced by ssRNA40. The induction of cytokine secretion was fully abolished in *TLR7* KO macrophage cells (**Fig. 5D**), indicating that these rRFs activate TLR7 to promote cytokine production.

**Figure 5.**
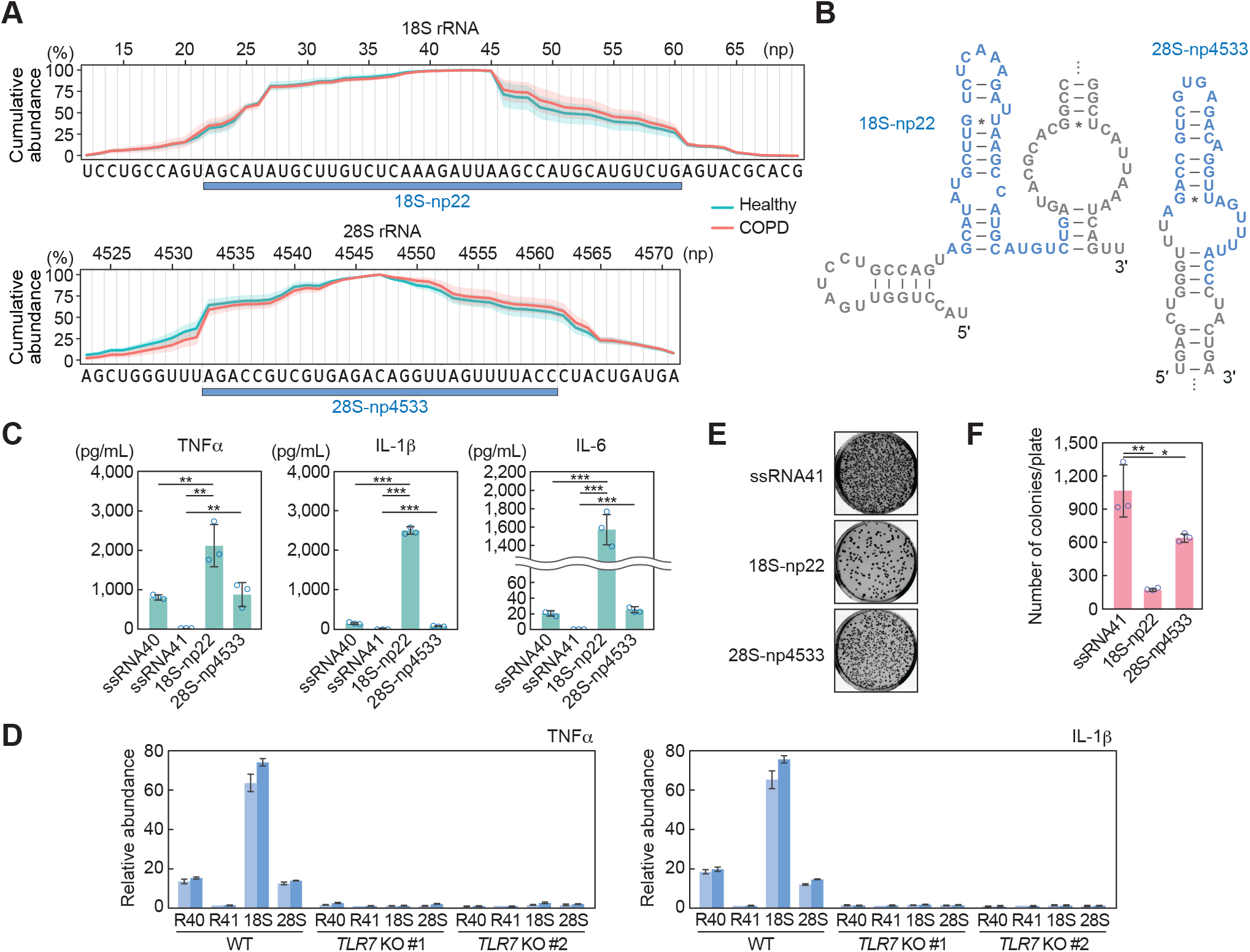
rRFs activate endosomal TLR in human macrophages. **(A)** Cumulative plot of the plasma sncRNA reads in the selected regions of 18S and 28S rRNAs.The regions generating two specific rRFs, 18S-np22 and 28S-np4533, are shown. The shaded regions indicate SD from four replicates. **(B)** Secondary structure of the rRFs. The targeted rRF plus 50-nt upstream and downstream region were used for prediction. **(C)** After DOTAP-mediated endosomal delivery of the indicated RNAs into macrophages, culture medium was subjected to measurement of concentration of the indicated cytokines. The error bars denote the SD of three independent experiments. \*\**p*<0.01; \*\*\**p*<0.001. **(D)** The DOTAP experiments were performed by using *TLR7* KO macrophages, followed by quantification of cytokine mRNAs by RT-qPCR. Two independent experiments were performed, which are shown as separated bars, while the error bars indicate the SD of qPCR replicates. **(E, F)** After DOTAP-mediated endosomal delivery of the indicated RNAs, macrophages were subjected to bacterial infection and invasion assay. Representative pictures of the plates with *E.coli* colonies (E) and bar graphs of the counted numbers of colonies (F) are shown. The error bars indicate the SD of three independent experiments. \**p*<0.05; \*\**p*<0.01.

We also performed a bacterial elimination assay, showing successful elimination of infected *E.coli* after the endosomal delivery of 18S-np22 and 28S-np4533 (**Fig. 5E and 5F**). 18S-np22 showed stronger activity than 28S-np4533, which is consistent with the more substantial cytokine secretion by 18S-np22 compared to 28S-np4533. These results suggest that the abundantly accumulated rRFs, as well as 5′-tRNA halves, can serve as immune-modulator molecules by stimulating endosomal TLR.

## Discussion

Circulating RNAs have emerged as novel biomarkers and functional molecules in diverse areas of disease studies. While previous investigations into circulating sncRNAs in the biofluids of COPD patients have primarily focused on miRNAs through the use of standard RNA-seq ^25–28^, it is important to note that the majority of extracellular sncRNAs contain either a cP or 3′-P ^24^ and are thus not captured by standard RNA-seq. Moreover, even a single cP-containing tRNA half, rRF, or mRF species can be significantly more abundant than total miRNAs in tissues ^30^ and EVs ^24^. This suggests that standard RNA-seq data contains critical biases by excluding the majority of circulating sncRNAs ^21–23^. In this study, by incorporating pre-treatment of plasma RNA with T4 PNK into sncRNA sequencing process, we obtained a comprehensive and accurate representation of sncRNAs in human plasma samples and discerned their differential profiles between healthy individuals and COPD patients. While we acknowledge certain limitations in the present study including the small sample size and the lack of the detailed clinical information, our studies provide the first sequence information of circulating sncRNAs in COPD patients and the basis for exploring unique species of sncRNAs as biomarkers in COPD diagnosis. These approaches, which can identify circulating sncRNAs that elude standard RNA-seq—such as cP-containing 5′-tRNA halves ^19, 34^ that are actually are predominant in circulating tRNA-derived sncRNAs—should be implemented in exRNA research for other diseases to advance our understanding of the molecular landscape and potential biomarkers. It is also important to note that post-transcriptional modifications of RNAs, which may impede reverse transcription, could introduce biases in sequencing results. Exploring potentially underrepresented sncRNAs due to modifications can be achieved by sncRNA sequencing procedures that include a step to remove some of the RT-hindering modifications ^47–49^.

Focusing on tRNA-derived sncRNAs, we demonstrated that the levels of 5′-ValCAC are significantly upregulated in COPD patients, making these molecules the most abundant circulating tRNA-derived sncRNAs. In contrast, in healthy individuals, we observed 5′-GlyGCC to be the most abundant tRNA-derived sncRNA. Of note, dimerization of the molecule has been shown to increase its stability ^50^, potentially contributing to its abundant accumulation in circulation. While the predominant existence of 5′-GlyGCC as a circulating sncRNA aligns with previous studies ^51^, the sequence detected in our study is longer, specifically a 34-nt molecule of 5′-GlyGCC. We assume that the lack of T4 PNK treatment in sncRNA sequencing could result in the selective detection of secondary cleaved fragments that contain a 3′-OH end.

Further research is required to explore potential molecular mechanisms governing the differential expression patterns of circulating sncRNAs. One potential mechanism underlying the increased levels of 5′-ValCAC in COPD patients could be enhanced anticodon cleavage activity for tRNA^ValCAC^. Enhanced RNA cleavage in COPD is suggested by the enrichment of rRFs with high GC content. The nucleotide composition analysis of the 3′-ends of 3′-tRNA halves may also support this, as the proportion of 3′-halves with an intact CCA-end was higher in COPD patients, suggesting an accelerated cleavage of mature tRNAs with CCA to produce 3′-halves. The enhanced RNA cleavage may also account for the observed positive correlation among 5′-half, 5′-tRF-S, and i-tRF-S in COPD. As 5′-ValCAC accumulates, its shorter derivatives are also produced in plasma. Furthermore, our nucleotide composition analyses suggest preferable cleavages occur between C and A or U and A in the generation of circulating rRFs and mRFs; consistent with this, the cleavage to produce 5′-ValCAC occurs between C and A. In contrast, tRNA^GlyGCC^ is likely not a preferred target for anticodon cleavage in COPD conditions, as 5′-GlyGCC levels were not upregulated, and instead, the proportion of i-tRF-5′ was significantly increased. Oxidative stress, identified as a major pathogenic driver in COPD ^52^, could induce these cleavages as evidenced by previous studies on stress-induced expression of tRNA halves, rRFs, and mRFs ^36, 53^. Alternatively, the specific upregulation of circulating sncRNAs could be achieved through stabilization after cleavage, such as by interacting with stabilizer proteins. YBX-1, a well-characterized transcription factor with extended functional roles in RNA biology, may play a role in the upregulation of 5′-halves, because YBX-1 binds to 5′-halves ^54, 55^ and plays a role in sncRNA packaging into EVs ^14^. These potential mechanisms, enhanced cleavage and stabilization, are expected not to be mutually exclusive. A recent study suggests that extracellular 5′- and 3′-tRNA halves can exist as nicked tRNA structures ^56^. Concurrently, other studies have shown distinct profiles and pathways for 5′- and 3′-tRNA halves within cells and during EV-packaging ^24, 57^. Distinct clustering patterns observed in PCA among tRNA-, rRNA-, and mRNA-derived sncRNAs suggest that each class of sncRNAs could be subjected to different regulatory mechanisms. Further research is required to investigate the potential molecular mechanisms that govern the differential expression patterns of circulating sncRNAs in COPD. While miRNAs are primarily associated with gene regulation, our results suggest that tRNA-and rRNA-derived sncRNAs may play an immunomodulatory role in COPD pathogenesis. Our previous experiments, in which human plasma was treated with a ribonuclease in the presence or absence of a detergent, suggest that the majority of plasma 5′-tRNA halves can be incorporated into EVs ^24, 41^. Given that the content of EVs is delivered to endosomes of the recipient cells, and that 5′-ValCAC can strongly activate TLR7 upon its delivery to endosome of macrophages, we speculate that 5′-ValCAC could function as a cytokine inducer by serving as the endogenous ligand for TLR7 in COPD pathogenesis. Although further research, particularly using an animal model, is necessary to examine this, our previous macrophage experiments have already shown that physiologically relevant amounts of 5′-HisGUG can potently induce cytokines ^24^, and the TLR7 stimulation activity of 5′-ValCAC is as strong as that of 5′-HisGUG ^41^. In this study, we further demonstrated that specific rRFs, which are upregulated in COPD patients, can induce cytokine production by activating endosomal TLR. Although the research on TLR7 in COPD patients has primarily focused on its association with viral infections and viral RNA has been considered its main ligand ^58–60^, we propose that circulating sncRNAs can act as strong ‘endogenous’ ssRNA ligands contributing to the observed cytokine production in COPD patients. Previous studies have shown increased cytokine production in COPD patients’ lungs upon stimulation by R848, a widely used TLR7/8 ligand ^61^. Therefore, TLR7 in these patients may exhibit heightened responsiveness to upregulated immune-active circulating sncRNAs. Importantly, we have demonstrated that 5′-GlyGCC does not activate TLR7 in macrophages ^41^. Selective mechanisms may exist in the production and/or stabilization of immune-active molecules in COPD patient plasma, while the degradation of non-active molecules may be promoted.

Numerous previous studies have identified subsets of miRNAs exhibiting distinct expression levels not only between healthy individuals and COPD patients but also in relation to smoking history ^62–66^. Considering the abundant presence of circulating tRNA- and rRNA-derived sncRNAs and their differential expression patterns between healthy individuals and COPD patients, further exploration of their expressional relationships with various pathological factors, including not only smoking status but also disease severity stages, is required. This investigation should be conducted with a larger sample size, which could designate these sncRNAs as biomarkers.

## Materials and Methods

### Ethical approval, human plasma samples, and RNA isolation

The Office of Human Research (OHR) of Thomas Jefferson University (TJU) approved our use of human plasma samples without personal information in accordance with all federal, institutional, and ethical guidelines. We obtained the de-identified human plasma samples from a biological specimen company, BioIVT. Human plasma samples were derived from healthy individuals and COPD patients (**Table S2**). RNA isolation from the plasma samples was performed as described previously ^24, 29^. First, 1,000 µL of plasma were centrifuged at 16,060 g for 5 min, then 800 µl of supernatant were subjected to RNA extraction using TRIzol LS (Invitrogen). The extracted RNAs were further subjected to column purification using the miRNeasy Mini Kit (Qiagen).

### Multiplex quantification of 5′-tRNA halves

The levels of 5′-tRNA halves were quantified by a multiplex TaqMan RT-qPCR method specifically developed for limited starting materials as described in our previous study ^29^. Briefly, 400 µL of plasma supernatant were mixed with 1 fmol of Spike-in control RNA (5′-CAGUGGUGGGCCAGAUGUAAACAAAUGAAUGUUCUUG-3′) and subjected to RNA extraction using TRIzol LS. The RNA pellet was resuspended in 8 µL of RNase-free water, and then 2 µL of RNA solution were used for T4 PNK treatment and 3′-adaptor ligation as described previously ^29^. Multiplex TaqMan RT-qPCR was performed using One Step PrimeScript RT-PCR Kit (Takara) on StepOne Plus Real-time PCR machine (Applied Biosystems). The Ct value of the Spike-in RNA was used for normalization. The sequences of targeted 5′-tRNA halves, primers, and TaqMan probes are shown in **Table S1 and S4**. The data were analyzed using the *R* package, *beeswarm*, and presented as a box plot.

### Pre-treatment of plasma RNA with T4 PNK and sncRNA sequencing

RNA extracted from 800 μL of plasma was treated with T4 PNK in the presence of 1 mM of ATP at 37°C for 40 min, followed by phenol/chloroform/isoamylalcohol extraction and ethanol precipitation. The recovered RNA was then subjected to cDNA amplification by using TruSeq Small RNA kit (Illumina). The amplified cDNAs were gel-purified and their quality and amount were assessed using Bioanalyzer High Sensitivity DNA chip (Agilent) and Qubit (Thermo Fisher Scientific). The cDNA libraries were pooled at 4 nM concentration and sequenced on Illumina NextSeq 500 at the MetaOmics Core Facility of the Sidney Kimmel Cancer Center at TJU.

### Data analysis

Bioinformatic analyses were performed as described previously ^30, 36^. In brief, we used the cutadapt tool (DOI: http://dx.doi.org/10.14806/ej.17.1.200) to remove the 3′-adaptor sequence. After selecting 18–50-nt reads, we used Bowtie2 for the sequential mappings ^67^. Reads were mapped to mature cytoplasmic tRNAs obtained from GtRNAdb (Release 18) ^38^, and then to mature rRNAs, to mRNAs, to the mitochondrial genome (NC_012920.1 sequence plus 22 mitochondrial tRNA sequences), and to the whole genome (GRCh37/hg19) ^30, 36^. Data analysis and visualization were carried out using *R* packages: *gplots* for heat map analysis; and *corrplot* and *devtools* for *Spearman* correlation analysis. The average values of correlation coefficient in healthy and COPD groups were calculated by *Spearman*’s Z-transformation and are presented as Z-values. The alignment of 5′-half, 5′-tRF-S, and i-tRF-5′ were visualized by *ggplot2*, *tidyverse*, and *dplyr*. Differential expression analysis for rRFs and mRFs was carried out using DESeq2. sncRNAs that showed less than one read throughout eight libraries were omitted for the analysis.

### *In vitro* RNA synthesis

The synthetic RNAs used in this study are shown in **Table S5**. They were synthesized by an *in vitro* reaction with T7 RNA polymerase (New England Biolabs) as described previously ^24, 40^. dsDNA templates were synthesized using PrimeSTAR GXL DNA Polymerase (Takara Bio) and the primers shown in **Table S4**. The synthesized RNAs were then gel-purified using denaturing PAGE with single-nucleotide resolution.

### Cell culture, endosomal delivery of RNA, and measurement of cytokine concentration

THP-1 human acute monocytic leukemia cells (American Type Culture Collection) were cultured in RPMI 1640 medium (Corning) with 10% FBS and differentiated into macrophages using phorbol 12-myristate 13-acetate (PMA; Sigma-Aldrich) as described previously ^24, 68^. *TLR7* KO THP-1 cell lines, whose TLR7 expression is completely depleted, were previously generated by using CRISPR/Cas9 approach ^24^. Before transfection, the cells were primed with 100 units/ml of interferon γ (Thermo Fisher Scientific) for 18-24 h ^69^. To deliver RNAs to endosomes, we used the cationic liposome 1,2-dioleoyloxy-3-trimethylammonium-propane (DOTAP, Sigma-Aldrich) as previously described ^24, 70, 71^. In brief, 230 pmol of synthetic RNAs were mixed with 60 µl of HBS buffer and 15 µl of DOTAP reagent and incubated for 15 min. The RNA-DOTAP solution was then added to 1 ml RPMI 1640 medium with 2% FBS, followed by incubation of the cells for 16 h. For quantification of cytokine mRNAs by RT-PCR, as in our previous study ^24^, total RNA was extracted from the cells using TRIsure (Bioline), treated with DNase I (Promega), and subjected to reverse transcription using RevertAid Reverse Transcriptase (Thermo Fisher Scientific) and a reverse primer. The synthesized cDNAs were then subjected to PCR using 2×qPCR Master Mix (Bioland Scientific) and forward and reverse primers, as described in our previous study ^24^. The abundance of the target mRNA was calculated as a ratio to GAPDH mRNA, and further normalized to the control experiment (transfection of ssRNA41). Cytokine concentrations of the cultured medium of macrophages were measured by Multiplexing LASER Bead Technology (Eve Technologies).

### Bacterial infection and invasion assay

After 16 h of DOTAP transfection of RNAs, THP1-derived macrophages (1 × 10^6^ cells) were plated on 6-well plates and incubated with *E. coli* [10 multiplicities of infection (MOI)] in RPMI 1640 (no antibiotics) for 60 min at 37°C. The cells were then washed with PBS three times and incubated with RPMI 1640 containing high concentration (3×) of penicillin-streptomycin (Thermo Fisher Scientific) at 37°C for 60 min. Subsequently, medium was replaced with RPMI 1640 containing normal concentration (1×) of penicillin-streptomycin, followed by further incubation at 37°C for 24 h. The cells were then washed and lysed with 0.5% Triton X-100. Intracellular bacteria were enumerated by plating on LB agar plates.

## Data availability

The sequence reads are publicly available from the NCBI Sequence Read Archive (under the BioProject: PRJNA985703).

## Author Contributions

The research was conceived by M.S., D.A.D., and Y.K. The experiments were designed by M.S. and Y.K., and were performed by M.S. and T.K. The data analysis was performed by M.S. The paper was written by M.S. and Y.K., with contributions from D.A.D. The funding support was provided by M.S., D.A.D., and Y.K.

## Acknowledgments

We are grateful to Justin Gumas (TJU) for helpful discussions and for the maintenance of Kirino Lab plasma collections, and to Takashi Hamasaki (Calpis America, Inc.) for the instructions on *R* packages, *tidyverse* and *dplyr*. This study was supported by the National Institutes of Health Grant (HL150560 to DD and YK; GM106047, AI151641, and AI168975 to YK) and American Lung Association Catalyst Award Grant (CA935540 to MS). This research utilized the MetaOmics Core Facility at Sidney Kimmel Cancer Center in TJU and was supported by the National Institutes of Health Grant (P30CA056036).

