## Supplementary material for "Immunoactive signatures of circulating tRNA- and rRNA-derived RNAs in chronic obstructive pulmonary disease": Figures S1-S5, Tables S1-S5

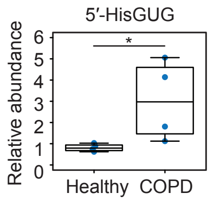


**Figure S1. The levels of 5′-HisGUG in plasma samples**

Plasma RNAs (Healthy: $n = 4$; COPD: $n = 4$) were subjected to TaqMan qPCR for specific quantification of 5′-HisGUG. Spike-in RNA was added during RNA extraction, and its levels were used for normalization. The value of one of the healthy samples was set as 1, and the relative values for the other samples are shown. *$p < 0.05$.

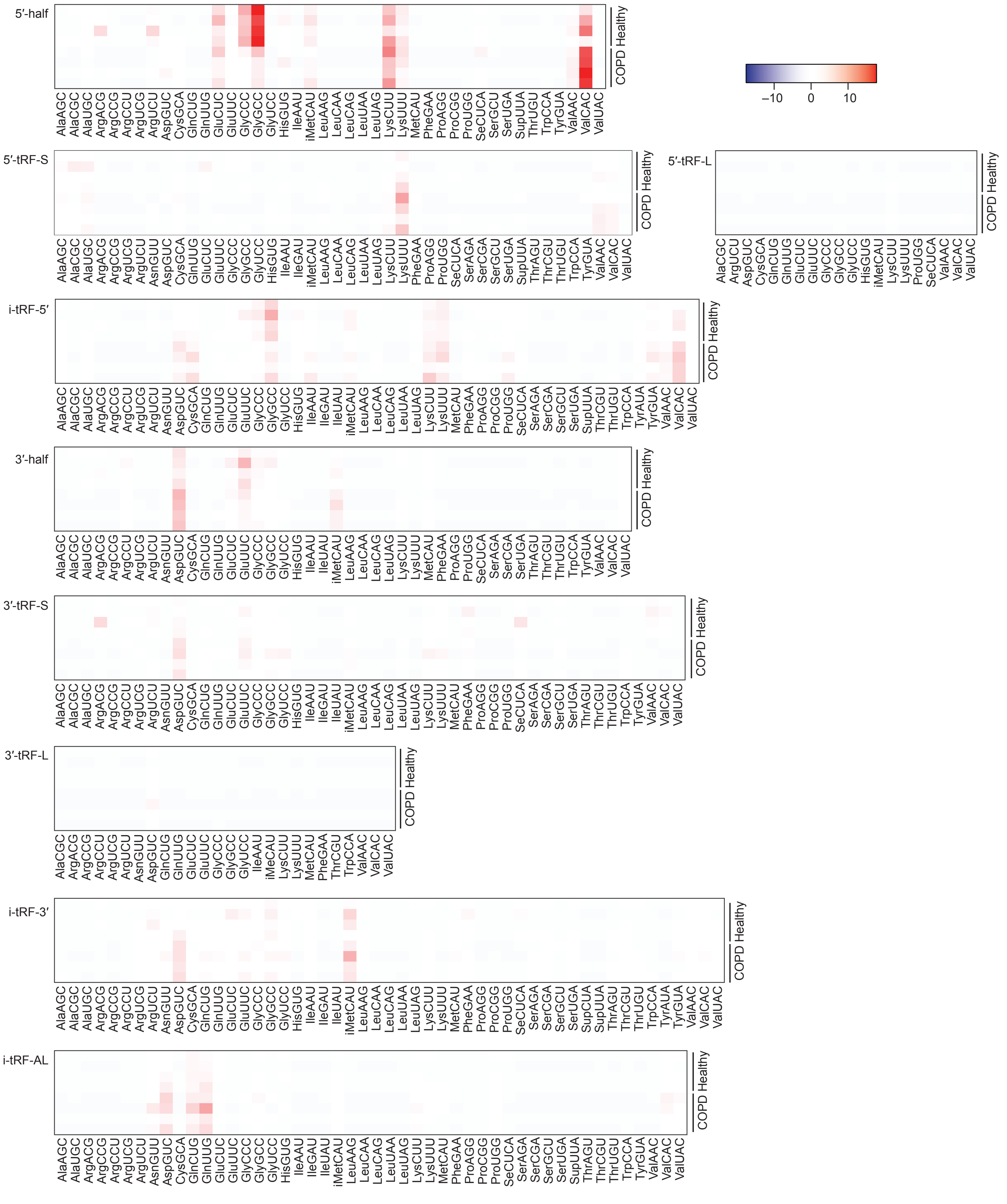


**Figure S2. Heatmap of each subclass of tRNA-derived sncRNAs**

The expression profiling was performed across all subclasses of tRNA-derived sncRNAs, and the results are presented in alphabetical order of tRNA isoacceptors.

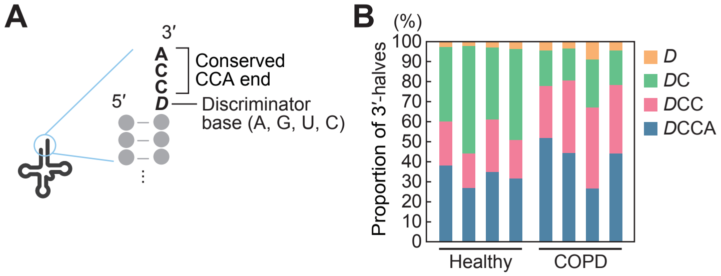


**Figure S3. Nucleotide analysis of the 3′-ends of 3′-halves**

(A) Schematic representation of tRNA termini. Mature tRNA commonly contains a 4-nt protruding 3′-end consisting of a discriminator base and conserved CCA trinucleotides.

(B) Proportions of each 3′-terminal variant of 3′-halves.

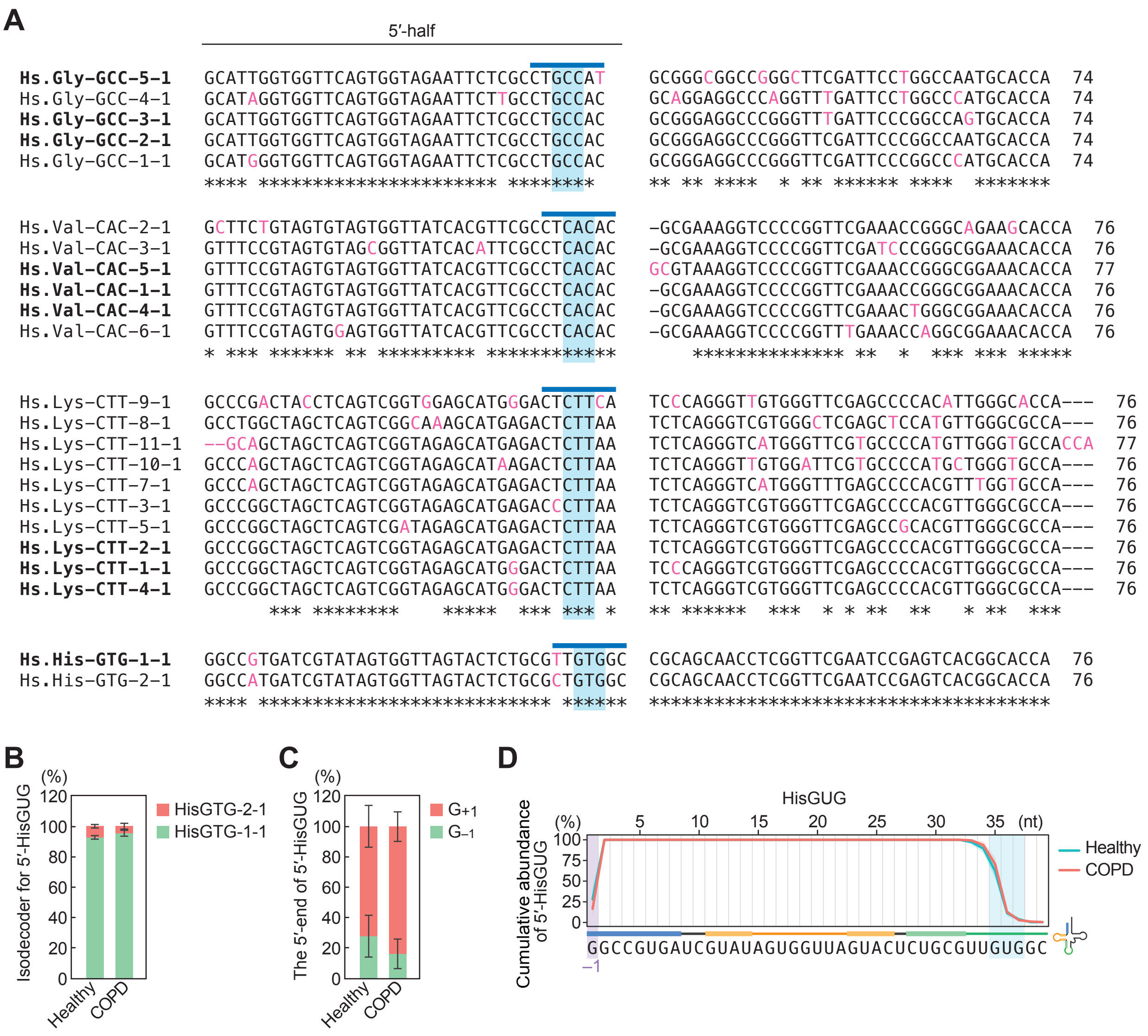


**Figure S4. Sequences of isodecoders and analysis of 5′-HisGUG**

(A) Sequence alignment of the four indicated tRNAs. The blue line indicates anticodon loop region, and light blue highlights the anticodon. The genes that produce 5′-halves are indicated in bold.

(B) Proportion of 5′-HisGUG for each isodecoder.

(C) Proportion of the 5′-terminal nucleotide of 5′-HisGUG.

(D) Cumulative line graph of 5′-HisGUG. The shaded region overlaying each line represents the S.D. from four replicates.

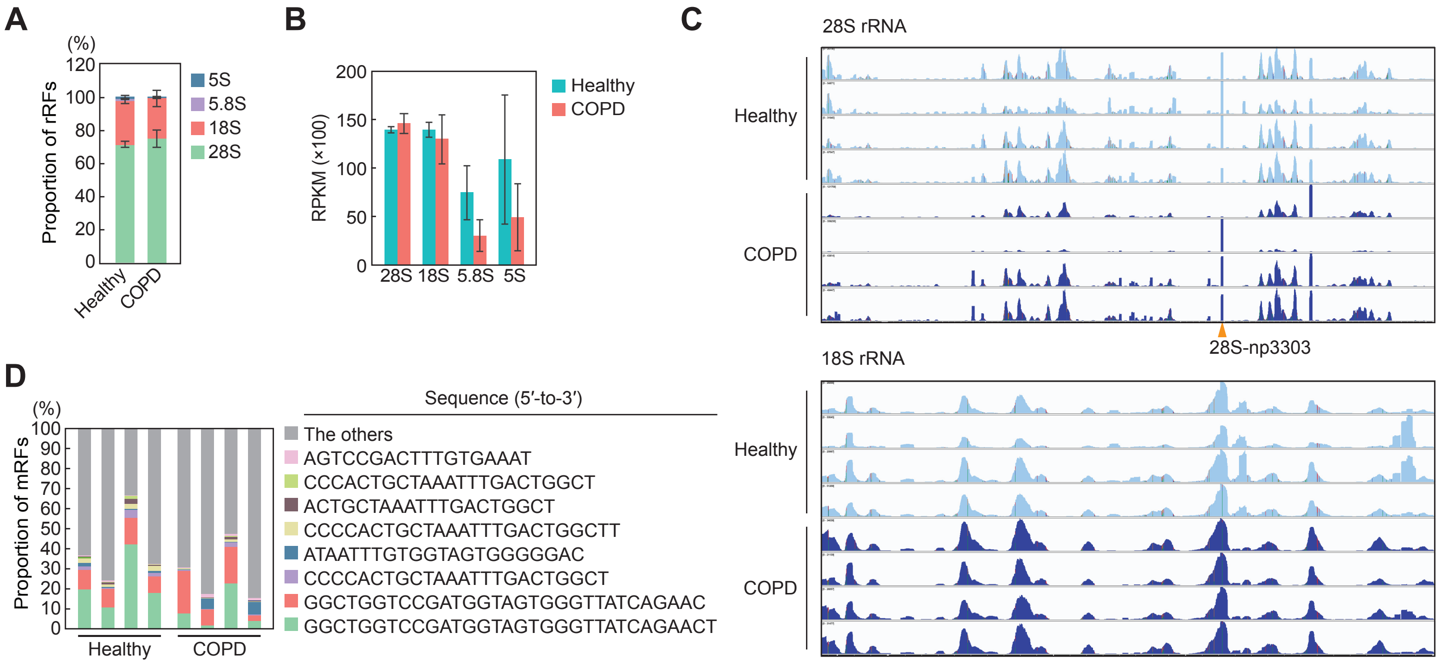


**Figure S5. Analysis of rRFs and mRFs**

(A) Proportion of rRFs for each rRNA gene. The error bar indicates the SD from four replicates.

(B) Abundance of rRFs is shown as RPKM of each rRNA gene. The error bar indicates the S.D. from four replicates.

(C) Read alignment of rRFs visualized by Integrative Genomics Viewer.

(D) Proportion of the 8 most abundant mRFs.

**Table S1. Unique ID for the sncRNAs investigated in this study**

| Name | Sequence (5′-to-3′) | License plate | tDRname |
|---|---|---|---|
| 5′-HisGUG | GCCGUGAUCGUAUAGUGGUUAGUACUCUGCGUUG | tRF-34-PW5SVP9N15WV2P | tDR-1:34-His-GTG-1 |
| 5′-GlyGCC | GCAUUGGUGGUUCAGUGGUAGAAUUCUCGCCUGC | tRF-34-PNR8YP9LON4VHM | tDR-1:35-Gly-GCC-2-M3 |
| 5′-ValCAC | GUUUCCGUAGUGUAGUGGUUAUCACGUUCGCCUC | tRF-34-79MP9P9NH57S15 | tDR-1:34-Val-CAC-1-M3 |
| 18S-np22 | AGCAUAUGCUUGUCUCAAAGAUUAAGCCAUGCAUGUCUG | rRF-39-FQII9720IOPQP4JP | (N/A) |
| 28S-np4533 | AGACCGUCGUGAGACAGGUUAGUUUUACC | rRF-29-F1SW5D1N1Z2V | (N/A) |

**Table S2. Plasma donor information**

| Category | ID | Exp # in Fig. 1H | BioIVT Lot ID | Experiments | 5'-Val value | Age | Gender | BMI | Race | Diagnosis (year) | Severity/ GOLD Stage | Smoking status | Smoking history | Smoking frequency | Medications | Previous Treatments |
|---|---|---|---|---|---|---|---|---|---|---|---|---|---|---|---|---|
| Healthy | 1 | Set 1 | HMN664977 | Val qPCR, RNA-seq | 0.537 | 56 | Male | 28 | Caucasian | Normal Donor | - | N/A | N/A | N/A | None | None or N/A |
| | 2 | Set 1 | HMN826748 | Val qPCR, RNA-seq, Gly qPCI | 1.433 | 61 | Female | 28 | Caucasian | Normal Donor | - | N/A | N/A | N/A | None | None or N/A |
| | 3 | Set 1 | HMN829012 | Val qPCR, RNA-seq, Gly qPCI | 1.006 | 64 | Male | 28 | Caucasian | Normal Donor | - | N/A | N/A | N/A | None | None or N/A |
| | 4 | Set 1 | HMN826749 | Val qPCR, Gly qPCR | 1.586 | 68 | Male | 32 | Caucasian | Normal donor | - | N/A | N/A | N/A | None | None or N/A |
| | 5 | Set 1 | HMN690294 | Val qPCR, RNA-seq, Gly qPCI | 0.818 | 70 | Male | 28 | Caucasian | Normal Donor | - | N/A | N/A | N/A | None | None or N/A |
| | 6 | Set 2 | 133-0488 | Val qPCR, Gly qPCR | 0.895 | 56 | Female | 31 | Caucasian | Normal Donor | - | N/A | N/A | N/A | None | None or N/A |
| | 7 | Set 2 | 133-0533 | Val qPCR, Gly qPCR | 2.167 | 56 | Female | 26 | Caucasian | Normal Donor | - | N/A | N/A | N/A | None | None or N/A |
| | 8 | Set 2 | HMN715572 | Val qPCR, Gly qPCR | 1.000 | 61 | Male | 34 | Caucasian | Normal Donor | - | N/A | N/A | N/A | None | None or N/A |
| | 9 | Set 2 | 133-0531 | Val qPCR, Gly qPCR | 0.555 | 67 | Female | 29 | Caucasian | Normal Donor | - | N/A | N/A | N/A | None | None or N/A |
| | 10 | Set 2 | HMN664941 | Val qPCR, Gly qPCR | 0.556 | 56 | Female | 30 | Caucasian | Normal Donor | - | N/A | N/A | N/A | None | None or N/A |
| COPD | 1 | Set 1 | HMN811067 | Val qPCR, RNA-seq, Gly qPCI | 1.394 | 66 | Female | 26 | Caucasian | COPD (N/A) | Mild | N/A | N/A | N/A | Albuterol sulfate HFA | None or N/A |
| | 2 | Set 1 | HMN811069 | Val qPCR, Gly qPCR | 1.365 | 63 | Male | 35 | Caucasian | COPD (N/A) | Severe | N/A | N/A | N/A | Trelegy, Prednisone, Naproxen | Albuterol |
| | 3 | Set 1 | HMN811070 | Val qPCR, Gly qPCR | 9.995 | 57 | Male | 32 | Caucasian | COPD (N/A) | Moderate | N/A | N/A | N/A | Advair, ProAir HFA, Albuterol and Ipratropium | Spiriva (2018-2019), Prednisone (11/27/2019-12/02/2019) |
| | 4 | Set 1 | HMN811071 | Val qPCR, RNA-seq, Gly qPCI | 3.824 | 53 | Female | 27 | Caucasian | COPD (N/A) | Stage 3 | N/A | N/A | N/A | Pulmicort, Flonase, Albuterol, Ventolin | None or N/A |
| | 5 | Set 1 | HMN811072 | Val qPCR, Gly qPCR | 1.022 | 69 | Male | 31 | Caucasian | COPD (N/A) | Severe, Stage 3 | N/A | N/A | N/A | Advair Diskus, Levofloxacin, Prednisone, ProAir, Trelegy Ellipta, Tudorza Pressai | None or N/A |
| | 6 | Set 1 | HMN811073 | Val qPCR, RNA-seq, Gly qPCI | 9.969 | 51 | Female | 33 | Caucasian | COPD (N/A) | Mild, Stage 1 | N/A | N/A | N/A | Ipratropium-Albuterol | None or N/A |
| | 7 | Set 1 | HMN811074 | Val qPCR, RNA-seq, Gly qPCI | 3.262 | 62 | Male | 27 | Caucasian | COPD (N/A) | Mild | N/A | N/A | N/A | Symbicort, Ventolin HFA | None or N/A |
| | 8 | Set 2 | HMN875976 | Val qPCR, Gly qPCR | 1.213 | 64 | Male | 33 | Caucasian | COPD (N/A) | N/A | N/A | N/A | N/A | None | Cardiac Stent Placement (2008) |
| | 9 | Set 2 | 259155A2 | Val qPCR, Gly qPCR | 3.801 | 57 | Male | 31 | Caucasian | COPD (1994) | N/A | Never Used | - | - | Prednisone | None or N/A |
| | 10 | Set 2 | 259157A3 | Val qPCR, Gly qPCR | 6.098 | 69 | Female | 30 | Caucasian | COPD (2000) | N/A | Current Use | For 54 years | 20 smoke(s)/day | Advair Diskus, Oxygen, Ipratropium bromide, Singulair | None or N/A |
| | 11 | Set 2 | 286170A1 | Val qPCR, Gly qPCR | 2.976 | 60 | Male | 31 | Caucasian | COPD (2006) | N/A | Previous Use | For 32 years | 10 smoke(s)/day | Albuterol, Asmanex | None or N/A |
| | 12 | Set 2 | 286193A2 | Val qPCR, Gly qPCR | 8.542 | 66 | Female | 29 | Caucasian | COPD (2000) | N/A | Previous Use | For 25 years | 20 smoke(s)/day | Theophylline, Spiriva, Advair, Fluocinonide ointment | None or N/A |
| | 13 | Set 2 | 338854A2 | Val qPCR, Gly qPCR | 1.833 | 60 | Male | 27 | Caucasian | COPD (2013) | N/A | Current Use | For >5 years | 5 smoke(s)/day | Symbicort, Salbutamol | None or N/A |
| | 14 | Set 2 | 338845A2 | Val qPCR, Gly qPCR | 1.727 | 61 | Male | 29 | Caucasian | COPD (2013) | N/A | Current Use | For >20 years | 10 smoke(s)/day | Fluticasone and salmeterol, Salbutamol | None or N/A |
| | 15 | Set 2 | 485043A1 | Val qPCR, Gly qPCR | 0.494 | 55 | Male | 23 | Caucasian | COPD (2015) | N/A | Previous Use | 25 years | 20 smoke(s)/day | Albuterol, Symbicort | None or N/A |

N/A: Not available

Healthy: *N* = 10 (average 61.5 ± 5.5), 50% female

COPD: *N* = 15 (average 60.9 ± 5.5), 33% female

**Supplemental Table 3. Proportion of mapped reads (%)**

| Group | Library | tRNA | rRNA | mRNA | Mitocondria transcript | Genome |
|---|---|---|---|---|---|---|
| Healthy | 1 | 1.677 | 59.816 | 3.866 | 0.794 | 33.847 |
| | 2 | 2.382 | 69.006 | 3.212 | 0.670 | 24.730 |
| | 3 | 1.480 | 51.871 | 10.619 | 0.876 | 35.153 |
| | 4 | 1.587 | 68.192 | 2.778 | 0.552 | 26.890 |
| COPD | 1 | 1.143 | 70.956 | 2.647 | 1.343 | 23.910 |
| | 2 | 0.589 | 61.756 | 2.904 | 0.097 | 34.654 |
| | 3 | 2.188 | 52.714 | 6.475 | 2.634 | 35.989 |
| | 4 | 1.103 | 49.072 | 3.829 | 0.292 | 45.704 |

**Table S4. Sequences of the oligos used in this study**

| Experiment | Target | Type | Sequence (5′-to-3′) |
|---|---|---|---|
| TaqMan qPCR | 5′-HisGUG | Forward primer | GCTCGCCGTGATCGTATAGT |
| | | TaqMan probe | /5HEX/TAGTACTCT/ZEN/GCGTTGGAACACTGCGTTTGC/3IABkFQ/ |
| | 5′-GlyGCC | Forward primer | GCATTGGTGGTTCAGTGGT |
| | | TaqMan probe | 6-FAM/ATTCTCGCC/ZEN/TGCGAACACTGCG/3IABkFQ/ |
| | 5′-ValCAC | Forward primer | GCTCGTTTCCGTAGTGTAGTGGT |
| | | TaqMan probe | /56-FAM/ACGTTCGCC/ZEN/TCGAACACTGCGTT/3IABkFQ/ |
| | R-Luc | Forward primer | CAGTGGTGGGCCAGATGT |
| | | TaqMan probe | /5HEX/TTCTTGGAA/ZEN/CACTGCGTTTG/3IABkFQ/ |
| *In vitro* transcription | 5′-ValCAC | Forward primer | CCTGCAGTAATACGACTCACTATAGGGAGACTACGGAAACCTGATGAGTCCGTGAGGAC |
| | | Reverse primer | mGmAGGCGAACGTGATAACCACTACACTACGGAAACGACGGTACCGGGTACCGTTTCGTCCTCACGGACT |
| | 18S-np22 | Forward primer | CCTGCAGTAATACGACTCACTATAGGGAGAAGCATATGCTCTGATGAGTCCGTGAGGAC |
| | | Reverse primer | mAmGACATGCATGGCTTAATCTTTGAGACAAGCATATGCTGACGGTACCGGGTACCGTTTCGTCCTCACGGACT |
| | 28S-np4533 | Forward primer | CCTGCAGTAATACGACTCACTATAGGGAGAACGACGGTCTCTGATGAGTCCGTGAGGAC |
| | | Reverse primer | mGmGTAAAACTAACCTGTCTCACGACGGTCTGACGGTACCGGGTACCGTTTCGTCCTCACGGACT |

**Table S5. Sequences of the synthetic RNAs used in this study**

| Name | Sequence (5′-to-3′) |
|---|---|
| 5′-ValCAC | GUUUCCGUAGUGUAGUGGUUAUCACGUUCGCCUC |
| 18S-np22 | AGCAUAUGCUUGUCUCAAAGAUUAAGCCAUGCAUGUCUG |
| 28S-np4533 | AGACCGUCGUGAGACAGGUUAGUUUUACC |
| ssRNA40 | GCCCGUCUGUUGUGUGACUC |
| ssRNA41 | GCCCGACAGAAGAGAGACAC |
